# Contribution of mobile elements to the uniqueness of human genome with more than 15,000 human-specific insertions

**DOI:** 10.1101/083295

**Authors:** Wanxiangfu Tang, Seyoung Mun, Adiya Joshi, Kyundong Han, Ping Liang

## Abstract

Mobile elements (MEs) collectively constituted to at least 51% of the human genome. Due to their past incremental accumulation and ongoing DNA transposition of members from certain subfamilies, MEs serve as a significant source for both inter- and intra-species genetic diversity during primate and human evolution. Since MEs can exert direct impact on gene function via a plethora of mechanism, it is believed that the ME-derived genetic diversity has contributed to the phenotypic differences between human and non-human primates, as well as among human populations and individuals. To define the specific contribution of MEs in making *Human sapiens* as a biologically unique species, we aim to compile a complete list of MEs that are only uniquely present in the human genome, i.e., human-specific MEs (HS-MEs).

By making use of the most recent reference genome sequences for human and many other primates and a unbiased more robust and integrative multi-way comparative genomic approach, we identified a total of 15,463 HS-MEs. This list of HS-MEs represents a 120% increase from prior studies with over 8,000 being newly identified as HS-MEs. Collectively, these ~15,000 HS-MEs have contributed to a total of 15 million base pair (Mbp) sequence increase through insertion, generation of target site duplications, and transductions, as well as a 0.5 Mbp sequence loss via insertion-mediated deletions, leading to a net total of 14.5 Mbp genome size increase. Other new observations made with these HS-MEs include: 1) identification of several additional ME subfamilies with significant transposition activities not visible with prior smaller datasets (e.g. L1HS, L1PA2, and HERV-K); 2) A clear similarity of the retrotransposition mechanism among L1, *Alu*s, and SVAs that is distinct from HERVs based on the pre-integration site sequence motifs; 3) Y-chromosome as a strikingly hot target for HS-MEs, particularly for LTRs, which showed an insertion rate 15 times higher than the genome average; 4) among the ME types, SVAs seem to show a very strong bias in inserting into existing SVAs. Among the HS-MEs, more than 8,000 elements were integrated into the vicinity of ~4900 unique genes, in regions including CDS, untranslated exon regions, promoters, and introns of protein coding genes, as well as promoters and exons of non-coding RNAs. In seven cases, MEs participate in protein coding. Furthermore, 1,213 HS-MEs contributed to a total of 3,124 experimentally identified binding sites for 146 of the 161 transcriptional factors in association with 622 genes. All these data suggest that these HS-MEs, despite being very young, already showed sufficient sign for their participation in gene function via regulation of transcription, splicing, and protein coding, with more potential for future participation.

In conclusion, our results demonstrate that the amount of MEs uniquely occurred in the human genome is much higher than previously known, and we predict that the same is true regarding their impact on human genome evolution and function. The comprehensive list of HS-MEs provides an important reference resource for studying the impact of DNA transposition in human genome evolution and gene function.

## Introduction

Like in most genomes of higher organisms, transposable elements or mobile element insertions (MEs hereafter) constitute a major component in the human genome. Reported percentages of MEs in the human genome ranges from 45 to 48.5% [1–3] and our analysis in this report showing as 52.1% based on the most recent reference genome sequences. This percentage is likely to increase slightly with further improvement of the genome sequences, especially those covering the constitutive heterochromatin regions, which tend to have a higher proportion of repetitive sequences than the genome on average, and with better tools for detecting old and more diverged MEs. In the human genome, MEs are mainly represented by retrotransposons, which propagate in a copy and paste fashion via transcribed RNAs as the intermediates, and they can be classified as the LTR (long terminal repeats) and non-LTR retrotransposons[2, 3]. The LTR group is characterized by the presence of LTRs at the two ends of internal sequences of viral origins and includes mainly the endogenous retrovirus (ERVs) or human ERVs (HERVs) for those in the human genome. These ERVs came from virus affecting germlines and integrated into the genomes during different stages of primate and human evolution, and together they constitute approximately 8% of the human genome[4]. The non-LTR retrotransposon group consists of several very distinct subtypes including short-interspersed elements (SINEs), Long interspersed elements (LINEs), and a chimeric type of elements (SINE-R/VNTR/*Alu* or SVA for short). They share the common properties of having a poly (A) 3’-end, possessing internal promoters, and retrotransposing by a L1-based target-primed reverse transcription (TPRT) mechanism [4, 5]. Among these, *Alu* elements, representing the relative young and most successful SINEs by number, have more than 1 million copies and contribute to ~13% of the human genome[4, 6]. L1s, representing the LINEs, have more than half million copies and make the largest contribution to the human genome by size (~17%), and members from this family with full coding capacity are responsible for transposing all non-LTR MEs [7–10]. Outside of SINEs and LINEs, SVAs emerged as the newest group of retrotransposons that are mainly found within the hominid group of primates with a relative small copy number in the human genome due to their short history [11, 12].

Despite initially considered to junk DNA implying that they had no function [13], research mostly from in last two decades has convincingly demonstrated that MEs make very significant contributions in shaping evolution of genomes and impacting gene function via a plethora of mechanisms. These mechanisms range from generation of insertional mutations and genomic instability, creation of new genes and splicing isoforms, and exon shuffling to alteration of gene expression and epigenetic regulation [14–20] [18, 21–24]. As endogenous insertional mutagens, MEs are also known to be responsible for causing certain genetic diseases in humans [25, 26].

More recently, MEs are also shown to contribute to the generation of tandem repeats and providing definitive regulatory function or regulatory potentials. It was shown that at least 23% of all tandem repeats, another type of repetitive elements apparently very different from transposable elements, is derived from transposable elements [27]. In a recent study, Ward and colleagues very interestingly demonstrated that under a heterologous regulatory environment, regulatory sites in MEs, including those specific to humans, can be activated to alter histone modifications and DNA methylation, as well as expression of nearby genes in both germline and somatic cells [28]. A profound implication of this result is that lineage- and species-specific MEs provide novel regulatory sites to the host genome, which can potentially offer regulation of nearby genes’ expression in a lineage- and species-specific manner, which ultimately leads to phenotypic differences. A very recent study nicely added such an example by showing that an ERV element is responsible for regulating innate immunity in humans by controlling the expression of adjacent IFN-induced genes [29].

Ongoing retrotransposition generates genetic diversity among species and among individuals within the same species [3, 7, 30, 31]. Therefore, analysis of species-specific MEs can help understand the roles of MEs in speciation and in species-specific phenotypes, with the study of HS-MEs attracting most of interest for many obvious reasons. In the human genome, certain members from L1, *Alu*, SVA, and HERV families are still active in retrotransposition. They are responsible for generating HS- MEs and MEs that are polymorphic among humans [10, 32, 33]. So far, a few studies have examined HS-MEs as being present in the human genome, but not in the orthologous regions of any other primate genomes [31, 34–36]. Among these, the study by Mills and colleagues, representing the most comprehensive analysis of species-specific at the genome-scale covering all types of MEs, identified a total of 7786 and 2933 MEs that are specific to human and chimpanzee, respectively [31]. This study was done with earlier versions of the human and chimpanzee genome sequences (GRHc35/hg19 and CGSC1.1/panTrol1.1), which contained more unsequenced regions and assembly errors, in particular for the chimpanzee genome, and with no other primate genome sequence available. The identification of species-specific MEs was based on a blastz-based pair-wise alignment of the human and chimpanzee genome sequences to detect deletions in one genome that correspond to MEs with detectable target site duplications (TSDs) in the other genome. Since then, the genome sequences of human and chimpanzee both have been improved with several major updates, and the genome sequences of several additional closely related primates have also become available [37–44]. These additional primate genomes can be useful in providing complementary information to chimpanzee genome sequences. Therefore, we reasoned that it worthwhile to re-exam the human-specific MEs based all available new genome sequences. In this study, we developed a more sensitive 4-way comparative genomic approach involving pair-wise comparison of the human genome with that of chimpanzee, gorilla, orangutan, and rhesus monkey. We identified a total of more than 15,000 HS-MEs, among which more than 8,500 were reported as HS-MEs for the first time. This dataset represents a major improvement over prior studies and permits us to provide a more complete and accurate picture about the recent DNA transpositions in the human genome.

## Materials and Methods

### Sources primate genomic sequences

The human genome sequences used in this study was the February 2009 human reference sequence (GRCh37/UCSC hg19). Four other non-human primate genome sequences included as outgroup species to the human genome included the chimpanzee genome (Feb. 2011, CSAC Pan_troglodytes-2.1.4/panTro4), orangutan genome (Jul. 2007, WUSTL version Pongo_albelii-2.0.2/ponAbe2), gorilla genome (Dec 2014, NCBI project 31265/gorGor4.1), and rhesus monkey genome (Oct. 2010 GCA_000230795.1/rheMac3). The gibbon genome was not included in this study, as the data was unavailable at the UCSC genome browser until very late stage of our study. All involved genome sequences in fasta format and the RepeatMasker annotation files were downloaded from the UCSC genomic website (http://genome.ucsc.edu) onto our local server for in-house analyses.

### Identification of human-specific mobile element sequences (HS-MEs)

**Pre-processing of human MEs:** The starting list of all mobile elements we used is the repetitive elements annotated in the human genome using RepeatMasker (http://hgdownload.soe.ucsc.edu/goldenPath/hg19/database/rmsk.txt.gz). Since RepeatMasker reports fragments of MEs interrupted by insertions (mostly by MEs and low complexity sequences), internal inversion or deletions as individual ME entries, we performed a pre-process to integrate these fragments back to ME sequences representing the original retrotransposition events. Doing so permits us to obtain a more accurate counting of the retrotransposition events and also importantly to collect the correct flanking sequences for identifying HS-MEs and TSDs. Briefly, we examined the MEs that are next to each other with distance up to 50 kb, and check their mapping positions in the repeat consensus sequence and orientation. If two neighboring MEs mapped to the same repeat consensus sequences in the neighboring non-overlapping regions and in the orientation, plus the presence of TSDs at the revised ME ends, we then treat these ME segments as one ME entry with the start and end positions adjusted accordingly. For LTR retroransposons, RepeatMasker reports the two LTRs and the internal viral sequences each as separate entries, (e.g., a full-length LTR is reported as three separate entries). For our purpose, we grouped all components of a full-length LTR as one entry.

**Identification of HS-MEs**: Our strategy for identifying HS-MEs is to examine the sequences at the insertion and its flanking regions for each of the MEs (after integration) annotated in the human reference genome and compare with the sequences of the orthologous regions in multiple closely related non-human primate genomes. If a ME is determined with confidence to be absent from the orthologous regions of all examined non-human primate genomes, then it is considered to be human-specific. Briefly, we used two tools, BLAT [45] and liftOver (http://genomes.ucsc.edu) for determining the orthologous sequences and the human-specific status of MEs using the aforementioned integrated RepeatMasker ME list as input. Only the MEs that are supported to be unique to human by both tools were included in the final list of HS-MEs. A flow chart of the method is shown in Fig. 1, and the detailed description is provided in the supplementary materials.

**Figure 1.**
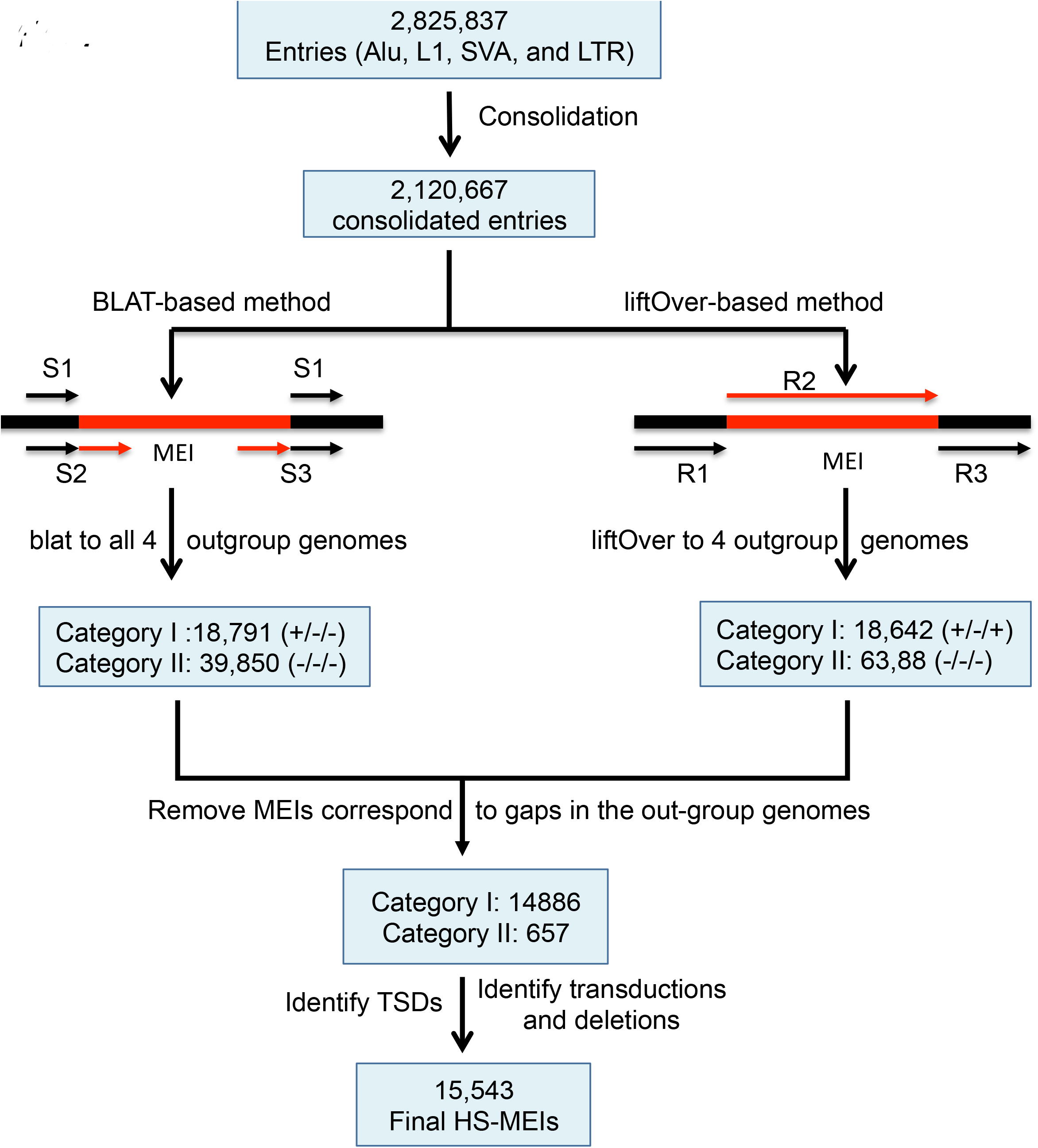
**Flow chart of identifying HS-MEIs**

### Validation of HS-MEs

To assess the accuracy of our HS-MEs both in terms of false-positive and false-negative errors, we performed manual inspection using the UCSC genome browser with a set of 100 randomly sampled HS-MEs, and performed further validations using the following three approaches.

<u>Method 1</u>: We compared our list against the previous HS-MEs list done by Mills et al in 2006, which was generated based on earlier versions of the human genome sequences using a different method (GRC35/UCSC hg17, May 2004).

<u>Method 2</u>: a set of reference MEs, which were detected as polymorphic as “deletions” by the 1000 Genome Project team [46, 47] and not yet covered by dbRIP, were collected and cross-matched with the list of HS-ME candidates.

<u>Method 3</u>: A total of randomly selected 16 HS-RE entries were sent to Kyudong Han’s research lab for PCR validation using a group of primate genome DNA samples (Details see supplementary materials and methods).

### Identification of TSDs and analysis of sequence motifs at the ME integration sites

The TSDs, as well as transduction and reotransposition insertion mediated-deletions (RIMDs) for all HS-MEs were identified using in-house Perl scripts incorporating the utility of the NCBI bl2seq and UCSC liftOver. It takes the human genomic sequence covering a HS-ME, plus flanking sequence on each side and aligns to the corresponding orthologous sequence from the chimpanzee genome (or the next closest genome with available orthologous sequences), which represents the pre-integration allele. In order to retrieve the pre-integration allele sequences, both flanking regions of HS-ME were liftOver’ed onto the out-group genomes. For typical category I HS-MEs, liftOver can find orthologous regions of the immediate flanking region. However, the immediate flanking of a HS-ME in human genome may not represent the pre-integration sequence due to possible transduction or RIMD. Therefore, our scripts liftOver’ed multiple subsequent blocks of flanking sequences retrieved in human genome onto the out-group genomes (100bp per block for up to 5Kb). The liftOver results were then grouped together to identify the shortest orthologous region containing the pre-integration site, which was aligned against the two human flanking sequences using blast. The overlapped region between the two aligned sections on the human sequence represents the TSD. For those with TSDs successfully identified, a 30-bp sequence centered at each insertion site in the predicted pre-integration alleles by removing the sequence of the ME and one copy of the TSDs from the ME alleles were extracted. The frequency of each of the 4 nucleotides at each position from 15 bp before and after the insertion site was calculated based on these pre-integration allele sequences, and plotted using the weblogo tool (http://weblogo.berkeley.edu/logo.cgi).

<u>Identification of ME-mediated transductions and deletions</u>: Entries with identified TSDs and extra sequences between the ME and either copy of the TSD are considered as potential candidates for ME-mediated transductions and were subject to further validation (see details in supplementary materials and methods). For entries without TSDs, if there are extra sequences at the pre-integration site in the out-group genomes, they were considered to be candidates for ME-mediated deletions in the human genome, which were subject to further validation.

### Analysis of HS-MEs in chromosome Y

The analysis of HS-MEs for chromosome Y required some special considerations for two reasons. First, authentic chromosome Y sequences are currently available only for chimpanzee and human, and they are not as complete as for autosomes[48]. Therefore, the analysis of HS-MEs can only be based on comparison with the chimpanzee sequences. Second, since the sequences for the pseudoautosomal regions (PARs) were basically copied from chromosome X, we excluded these from the analysis.

### Analysis of HS-ME distribution patterns in the human genome

The distribution patterns of HS-MEs in the human genome were examined by the density among different chromosomes and in the context of GC content and different regions of genes. For calculating GC content, 1500 bp sequences from both upstream and downstream of the ME insertions were used. These analyses were performed for each main type of MEs separately. The genomic coordinates of genes down to individual exons and coding regions were based on ENCODE[49] and NCBI RefSeq annotation[50] provided in the UCSC Genome Browser. The entire genome was divided into a non-redundant list of categorized regions in gene context as CDS, non-coding RNA, 5’-UTR, 3’-UTR, promoter (1kb), intron, and intergenic regions using an in-house perl script. This order of different genic region categories was used to set the priority from high to low in handling overlapping regions between different splice forms of the same gene or different genes. For example, if the same region is the coding region for one transcript and is an intron region for another transcript or for another gene, then this region would be categorized as coding region. For identifying HS-MEs overlapping with known transcriptional factor binding sites, we used the ENCODE Transcription Factor ChIP-seq contained in the wgEncodeRegTfbsClusteredV3.bed.gz available from the UCSC Genome Browser site.

## Result

### 1. A revised list of MEs in the human genome and a summary of HS-MEs

Our analysis of HS-MEs was based on the RepeatMasker annotation of repetitive elements in GRCh37/UCSC hg19 (February 2009), which were provided in the UCSC Genome Browser. To improve the accuracy in identifying HS-MEs and the TSDs and in calculating the rate of retrotransposition, we performed integration of fragmented MEs back to represent original ME events as detailed in the method section. This integration led to a reduction of almost one million (971,333) in ME counts from the 5,205,837 RepeatMasker ME entries in NCBI37/hg19 to 4,234,506 (Table S1). In the meantime, it resulted an increase of full-length MEs by 21% (data not shown). The number of annotated MEs showed a consistent increase in newer versions of the human reference genomes, especially among the earlier versions, mainly due to improved coverage of sequenced regions. The proportion of MEs in the genome increased from the earlier 48.8%[3] to 52.1% in the latest version (Table S1).

Using a multi-way comparative genomics approach involving comparison of human genome sequences to that of 4 other closely related primates including chimpanzee, gorilla, orangutan, and rhesus monkey, we identified a total of 15,463 human-specific HS-MEs. These HS-MEs consist of 9,764 *Alu*s, 3,710 L1s, 1,486 SVAs, and 503 HERVs by ME types, and 14,822 category I and 641 category II entries by the presence/absence of flanking sequence in the non-human primate genomes (Table 1). The complete list of HS-MEs is provided in supplementary file1 and in the dbRIP database. Among the HS-MEs, a total of 8,519 MEs were reported as HS-MEs for the first time, while 7,024 entries were shared with the previously reported HS-MEs[51] (Table S2).

**Table 1:** Summary of HS-MEIs

| MEI type | RM count | Integrated count | HS-MEIs |  |  |  |
| --- | --- | --- | --- | --- | --- | --- |
|  |  |  | Category I | Category II | Total | HS% |
| **L1** | 938,568 | 542,746 | 3,474 | 236 | 3,710 | 0.7 |
| **Alu** | 1,175,329 | 1,077,671 | 9,526 | 238 | 9,764 | 0.9 |
| **SVA** | 3,608 | 3,298 | 1,408 | 78 | 1,486 | 45.1 |
| **LTR** | 708,332 | 496,952 | 414 | 89 | 503 | 0.1 |
| **Total** | 2,825,837 | 2,120,667 | 14,822 | 641 | 15,463 | 0.7 |

### 2. Validation of HS-MEs

Due to the complexity of the task in identifying species-specific MEs as further discussed later, data generated by computational methods are very likely to have both false positives and false negatives. While it is not feasible for us to experimentally validate all 15,543 HS-MEs, mainly due to the extremely large number, we managed to validate the accuracy of our HS-ME list in three ways. First, we used the second half of the known polymorphic MEs documented in dbRIP [52] as a test dataset to check the rate of false negatives (i.e., sensitivity). Here, we reason that polymorphic MEs represent most recent insertions into the human genomes, and thus should be mostly human-specific. By cross-checking the HS-MEs with the 3110 polymorphic MEs (2331 from dbRIP and 779 from the 1000 genome project), which are included in the reference genome, we obtained an excellent rate of sensitivity of 95.5% (2972/3110).

Second, we cross-checked the HS-MEs with the 7,786 previously reported HS- MEs by Mills and colleagues [31], and we obtained 6,873 entries as shared. Among the 913 entries not on our list, we found that 787 actually represented false positives from the previous study and 8 were absent in the new version of the genome sequences, 47 entries for ME types not included in our input list, and 71 entries represent false negatives in our list (these were added to the final list of HS-MEs). This converts to a sensitivity of 98.9% (6,873/6,944) for our method. Since both the polymorphic MEs and those reported by Mills et al [31] are mostly from non-repetitive regions, which are less susceptible to errors in sequencing and computational analysis, the high ratio of overlapping is expected, and these were mostly identified as category I HS-MEs in our analysis as with matching orthologous sequences flanking the MEs in one or more non-primate genomes. We further compared our list with the human-specific HERVK list published by Shin and colleagues [36]. Among the 28 ME_IN entries, 26 of them overlap with our list while the other entries were not included in the final list due to low confidence.

Third, we performed validation using PCR, which is the gold stand for ascertaining MEs. Among the 16 loci we randomly selected, for which PCR was feasible for primer specificity and PCR product length, we were able to validate 92.3% (12/13), giving us a 7% of false positive rate. The entry failed to be validated in fact represents a chimpanzee-specific *Alu*-recombination mediated deletion, thus a chimeric *Alu* in the human genome, which was not handled by our method. Representative PCR results can be seen in Fig. S1 and full details of all examined loci are provided in Table S3.

Overall, combining the results from the above 4 types of validation analyses, the sensitivity and specificity of our HS-MEs is 95.4% (average of 98.1 and 92.3%) and 97%, respectively.

### Main sources for newly identified HS-MEs

We were interested to know the sources for these 8,469 novel HS-MEs. As shown in Table 2, the major contributors of novel HS-MEs are the MEs inserted into other MEs (3,089), a multiple factor group (“Others”) which include false negatives from prior study, less complete chimpanzee genome sequence, etc. (2,543), followed by non-canonical cases which involve transductions and insertion-mediated deletion or lack of TSDs (3,043). These 3 sources collectively contributed to 7,057 (83%) unique novel HS-MEs. Among the remaining 1,412 entries, consolidation of fragmented MEs contributed 830 entries and the use of more primate genome sequences beyond the chimpanzee genome added 526 entries, while the newer human genome sequences (from hg17 to hg19) added a modest of 56 entries.

**Table 2:** Sources for novel HS-MEIs (each factor considered by itself)

| **MEI type** | **Extra in hg19 (vs. hg17)** | **Entries consolidated from fragments** | **MEIs in MEIs** | **3 more primate genomes** | **Transduction/RIMD/no TSD** |
| --- | --- | --- | --- | --- | --- |
| **L1** | 10 | 555 | 739 | 254 | 1226 |
| **Alu** | 36 | 31 | 2099 | 541 | 1232 |
| **SVA** | 5 | 109 | 200 | 49 | 296 |
| **LTR** | 5 | 135 | 124 | 37 | 289 |
| **Total** | 56 | 830 | 3162 | 881 | 3043 |

### HS-MEs contributed to a 14 Mbp net size increase in the human genome

ME insertions can lead to genome size increase via insertion of MEs, generation of TSDs, and transductions, and they can also reduce genome size via retrotransposition insertion-mediated deletion (RIMD) of flanking sequences. As shown in Table 3, each type of MEs impacted the genome size using all 4 mechanisms and all made a net increase to the genome size. Collectively, all HS-MEs contributed to a total of 14.4Mb net genome size increase. Among the four types of MEs, L1s made the largest net increase (~8.2 Mb), followed by *Alu*s (~3.0 Mb), SVAs (~2.4Mb), and HERVs (~0.7Mb). HS-L1s contributed to the largest size changes using all mechanisms except for TSDs, and this is expected because of their large insert sizes and high copy number. Also as expected, HS-*Alu*s contributed to the largest size increase via TSDs by having the largest number. However, it is interesting to observe that L1s also contributed to the most transductions and RIMDs (Table 3). Despite having low retrotransposition activity, LTRs did contribute ~750kb size increase to the genome with 503 human-specific copies. Examples of HS-MEs carrying transductions and RIMDs can be seen in Fig. S2.

**Table 3:** Impact of HS-MEIs on genome size (Kb)

| Table 3: Impact of HS-MEIs on genome size (Kb) |  |  |  |  |  |
| --- | --- | --- | --- | --- | --- |
| MEI type | MEI insertions | TSDs | Transducitons | RIMDs | Total |
| **L1** | 7,860.0 | 36.4 | 444.8 | -144.7 | 8,196.5 |
| **Alu** | 2,833.8 | 118.7 | 155.2 | -64.5 | 3,043.3 |
| **SVA** | 2,140.9 | 19.2 | 272.7 | -14.0 | 2,418.8 |
| **LTR** | 713.9 | 1.8 | 89.6 | -46.5 | 758.9 |
| **Total** | **13,548.6** | **176.1** | **962.4** | **-269.7** | **14,417.5** |

\*RIMD, rertotransposition insertion-mediated deletion

### Recent retrotransposition activity level of MEs in the human genome

The comprehensive list of HS-MEs provided us an opportunity to re-examine more accurately the activities of MEs during human evolution, specifically since separation from chimpanzees. As shown in Table S4, while many observations were similar with what we learned from prior studies based mostly on younger polymorphic MEs ([3, 10, 31, 32]). These include that *Alu*, L1, SVA, and HERVs are the only four types of MEs with retrotransposition during human evolution, and *Alu*Ya5 and *Alu*Yb8/9, L1HS, SVA_E/D, and HERV_K are the most active subfamilies within the families of *Alu*, L1, SVA, and LTR, respectively. There are also several notable differences between the profiles of HS-MEs and polymorphic MEs. First, the ratio of HS-MEs in relation to all MEs in the active subfamilies were much higher (10 times or more) than the corresponding ratios of polymorphic MEs [10, 32] (Table S4). This might be a result of several factors, including a longer evolutionary span covered by HS-MEs, a likely decreasing retrotransposition during more recent human evolution, and an incomplete list of polymorphic MEs. Second, a few subfamilies extra to those represented by polymorphic MEs were seen among HS-MEs, including the *Alu*Yf and *Alu*Yk subfamilies from the *Alu* family, the L1P4 and L1M subfamilies from the L1 family, the SVA_A subfamily, and the ERV1 subfamilies from LTRs (Table S4). We reason that this is also likely due to the longer evolution span covered by HS-MEs and also likely that retrotransposition activity levels of these subfamilies have dropped to an undetectable level in the current human genomes.

### Patterns of MEs and HS-MEs in the repetitive regions

With our HS-MEs covering those in repetitive regions for the first time, we were interested in knowing whether there were any biases among the MEs both as insertion targets and as sources of insertions. We first examined the percentage of MEs inserting into other MEs for each ME type for all MEs and HS-MEs separately. For all MEs, there seems to be an increasing trend from LINEs (18.2%), DNA transposons (23%), LTRs (30.6%), SINEs (31.6%), and SVAs (44.5%) with the ratio of SVAs being more than doubled of that for LINE (Table S5a). We think these numbers reflect the relative overall age of ME types based on the amount of MEs existing in the genome during the span of propagation for the specific type of MEs. When only HS-MEs were considered (Table S5b), the ratio of those inserting into other MEs, showed a similar trend (with DNA transposons absent due to lack of human-specific members), but the degree of differences are smaller than for all MEs. This is expected since HS-MEs from all families share the approximately same time span during the human genome evolution, i.e. all types of HS-MEs had the same amount of MEs in the genome. Nevertheless, significant differences among the ME types are still observed, among which *Alu*s have the highest percentage of entries inserted into MEs (46%), followed by SVAs (42.5), and LTRs (39.1%), and L1 still have the lowest rate (35.8%) (Table S5b). Since HS-MEs from all families share the same time span during the human genome evolution, their ratios into MEs should provide a more accurate reflection of the preferences for MEs among ME types.

For MEs as the insertion targets of HS-MEs, we calculated the frequency of HS- MEs as the number of HS-MEs per million base pair of host ME sequences for each type of target MEs and HS-MEs (Table S5c). Among the ME types as targets, the HS-ME density follows a decreasing trend from high to low among LINEs (7.3), DNA transposons (5.0), LTRs (4.0), SINEs (1.8), reflecting their overall age in the genome from old to new. Very interestingly, SVA seems to be the exception to this trend as having the highest density of HS-MEs (13) despite being the youngest ME type (Table S5c). By HS-ME types, *Alu*s seem to be more frequently inserting into MEs than all other ME types except for SVAs as the targets. Remarkably, SVAs seem to show an unusually high self-preference. Specifically, HS-SVAs are inserted into SVAs at a frequency that is more than 18 times higher than into any other ME types (12.7 vs. 0.7), and SVAs as the targets received HS-SVAs at a frequency that is more than 60 times higher than other MEs (12.7 vs. 0.2; Table S5c). The mechanism for this unusual bias of SVAs inserting into SVAs and the functional implication are to be investigated in future studies.

### The characteristics of HS-MEs

To study the characteristics of HS-MEs, we examined the characteristics of HS- MEs, including the patterns of TSDs, integration site sequence motifs, distribution among chromosomes, and the GC content.

For distribution among chromosomes, we measured the HS-ME density (the number of HS-MEs per million base pairs of non-gaped chromosome sequences) and the ratio of HS-MEs among the same type of MEs in the chromosomes. As shown in Fig. 2 and Table S6, with all types of HS-MEs combined, the HS-ME density varies across chromosomes with Y chromosome showing the highest density (33.7 copies/Mb), more than 4 times higher than the genome average (7.3 copies/Mb) and more than 6 times higher than the lowest chromosome (chr22 at 5.0 copies/Mb). The distribution pattern is different among the 4 major types of MEs (Fig. 2). HS-*Alu*s showed a more or less homogenous distribution among the autosomes. However, Y chromosome also showed the highest density among all chromosomes, being more than double of that for the chromosome with the lowest. For L1s, the relative densities among chromosomes were more or less similar to *Alu*s for autosomes, while the density in Y chromosome is more than 3 times higher than the genome average. SVAs showed relatively similar densities among autosomes, except for chromosome 19, which has a density that is 2 times higher than the genome average, but it is extremely low (10 times lower than the genome average) in Y chromosome. LTRs showed the most biased distribution among chromosomes, with very low densities for most chromosomes, but with high, higher, and extremely high density in chromosome 19, 1, and Y, respectively. The HS-LTR density in Y chromosome (25.7 copies/100Mbp) is more than 15 times higher than the genome average (1.6 copies/100Mbp) and 30 times higher that of the chromosomes 10 and 13 (2.7 copies/100Mbp) (Table S6). Therefore, Y chromosome seems to be a hot target for *Alu*s, L1s, and LTRs with LTRs showing the highest preference over the rest of the genome, but it is the least preferred target for SVAs. Among other chromosomes, chromosome 19 stands out as having the highest density of all MEs, genes, HS-SVAs, and HS-LTRs, while chromosome 22 seems to have the least HS-*Alu*s, and HS-L1s. Correlation analysis showed a strong positive correlation of HS-SVA density with that of genes and with that of all MEs (R=0.9 in both cases), while the densities of HS-*Alu*s and HS-L1s showed a moderate positive correlation (R=0.6), and HS-L1s showed a moderate negative correlation with gene density (R=-0.4) (data not shown).

**Figure 2.**
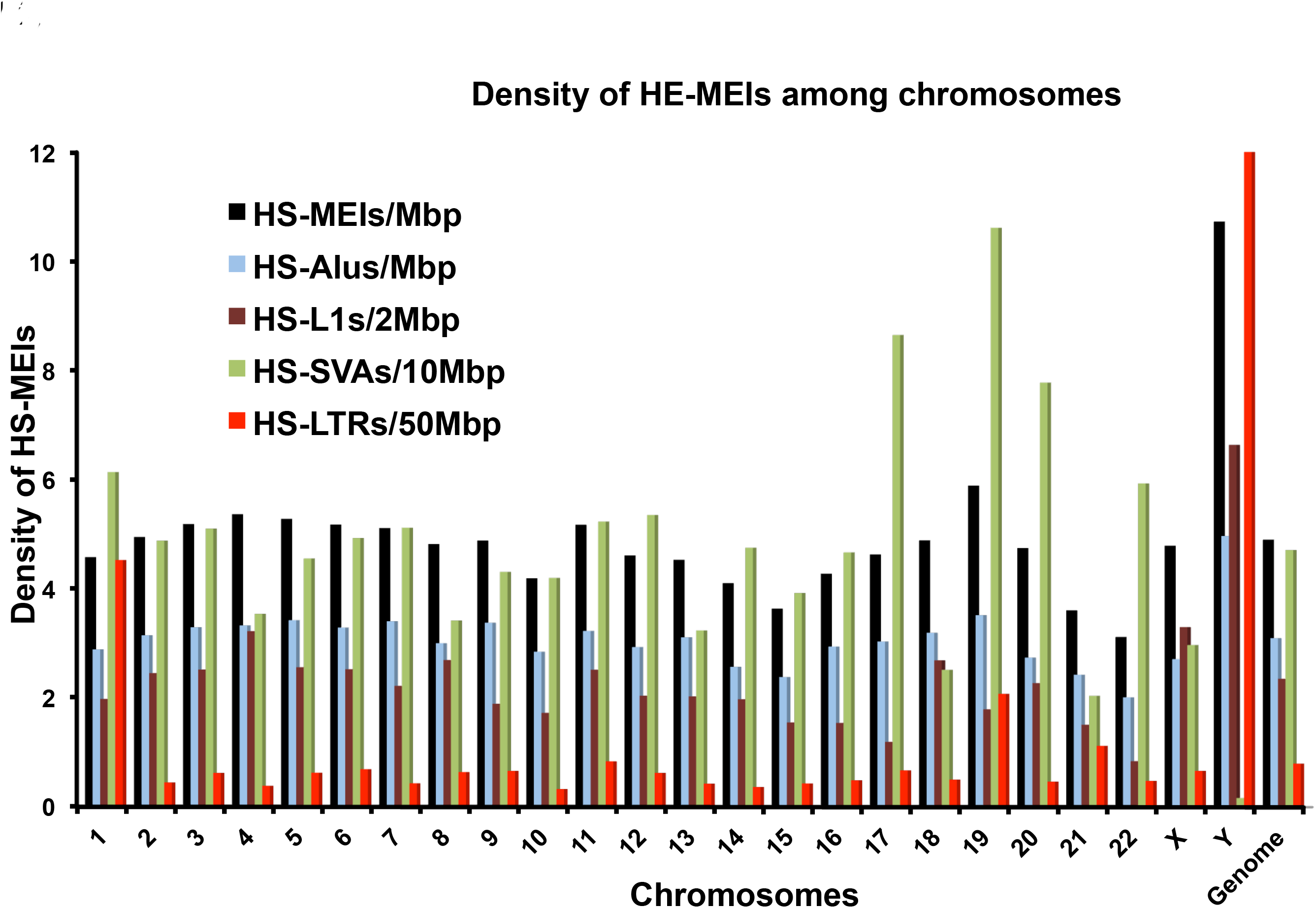
Genome distribution of HS-MEIs.

We calculated and compared the GC content for each ME type by using 1500 bp sequences both upstream and downstream of the ME insertions. Our data agrees with the pattern observed before that L1s tend to insert into GC-poor region and *Alu*s tend to insert into GC-rich regions[53], with the average GC content for being 42.1% and 39.0% for *Alu*s and L1, respectively in our data (data not shown). While HS-L1s are still inserting into GC-poor regions (37.8%), HS-*Alu*s tend to insertion into GC-poor region (39.4%) instead of GC-rich region. Both HS-SVAs and HS-LTRs tend to insert into GC-rich regions similar to their older, shared counterparts (Fig. S3).

We examined the sequence motifs at the integration sites for each ME type. As demonstrated by the sequence logos in Fig. 3A, *Alu*s, L1s, and SVAs showed an identical core sequence motif of “TT/AAAA”, confirming that all non-LTR retrotransposition events are driven by the TPRP mechanisms [54] [32] [55] [11]. The data also clearly show the distinction in site preference between LTR- and non-LTR retrotransposons with LTRs showing basically no recognizable target site motif, an observation not reported before despite their reported site preferences [56].

While the insertion site of MEs is determined by the involved endonucleases, we reason that the length of TSDs might reflect characteristics of different types of MEs beyond the involved endonucleases. As shown in Fig. 3B, all three non-LTR ME types showed a more or less similar distribution pattern peaking at the 15 bp. In the meantime, minor differences in the detailed shapes of the length distribution curves are also notable. For example, SVAs had a narrower peak, while *Alu*s showed a flatter peak covering 14 to 16 bp. Interestingly, L1s showed a secondary peak at 8 bp in addition to the main peak at 15bp, while LTRs also showed a bi-modular pattern with a main peak at 8 bp and a secondary peak at 6 bp. The latter is the feature for most ERVs. Again, these data clearly indicate the differences of the retrotransposition mechanisms used by LTR and non-LTR insertions, as well as some minor differences among different types of non-LTR insertions, which might reflect the impact of different characteristics of the ME sequences (lengths, GC composition, etc.) on the generation of the second nick, which determines the length of TSDs.

### HS-MEs’ Impact on genes and regulatory elements

To predict the functional impact of HS-MEs, we analyzed their genomic locations in gene context using the most updated version (Release 23, July 2015) of gene annotation data from the GENCODE project [49] combined with those in NCBI RefGene [50]. A total of 8,001 HS-MEs, representing more than half of all HS-MEs, are located inside or in the 1kb promoter regions of genes for protein coding, non-coding RNAs, as well as transcribed pseudogenes, representing a total of 4,713 unique genes/transcripts. As summarized in Table 4, while majority of these HS-MEs are located inside introns, a significant number (284) of them also directly participated in the transcriptomes as part of exon regions of protein coding genes, non-coding RNAs, and transcribed pseudogenes. Among those involved in the exons of protein-coding genes, 7 also contribute to protein coding. In all of these 7 cases, the involved transcripts represent rare splice forms that have not been documented in the NCBI’s refSeq genes, but reported in the GENCODE dataset.

**Table 4.** HS-MEIs in genic regions

| **Gene region** | **Protein coding** | **Non-coding RNA** | **Transcribed pseudogenes** | **Total** | **ratio (%)** |
| --- | --- | --- | --- | --- | --- |
| 1kb Promoter | 59 | 88 | 4 | 151 | 2.0 |
| CDS exon | 7 | NA | NA | 7 | 0.1 |
| non-coding exons | 144 | 130 | 3 | 277 | 3.6 |
| Intron | 5630 | 1532 | 118 | 7280 | 94.4 |
| **Total** | **5840** | **1750** | **125** | **7715** | **100.0** |

We also examined the contribution of HS-MEs in regulatory elements, specifically the ENCODE CHIP-seq transcriptional factor binding sites (TFBSs) for 161 factors[57]. Among the HS-MEs, 1,213 elements are overlapped with a total of 3,124 TFBS for 146 of the 161 transcriptional factors that are associated with 622 genes. Interestingly, by ME type, HS-LTRs have the highest ratio (15%) contributing to TFBS, followed by HS- SVAs (12%), which are much higher than the ratio for LINEs (8.1%) and SINEs (6.9%), although SINEs contribute to the largest number of TFBS (1,843), followed by LINEs (642), SVAs (444), and LTRs (195).

Our data suggest that these young HS-MEs have started making their ways in participating in the protein coding, transcripts, and regulation of splicing and transcription.

## Deposition of HS-ME data in dbRIP

In order for other researchers to easily access the HS-ME list, we deposited the data into the dbRIP database under a study ID of 2015-01. In dbRIP, the HS-MEs can be visualized in the same as the polymorphic MEs, i.e. in the UCSC genome browser along with other available data tracks or in detailed data page. To distinguish the HS- MEs from polymorphic MEs, all dbRIP IDs of the HS-MEs carry a letter of “h” at the end. The HS-ME data is also available for downloading at the dbRIP website (dbrip.org).

## Discussions

In this study, we aimed to provide an accurate compilation of mobile elements that are uniquely present in the human genomes, as a starting point for providing a comprehensive assessment about the impacts of mobile elements on human biology. By taking the advantages of the much-improved reference genome sequences for humans and the closely related non-human primates, including the mobile elements inside repetitive sequences, and utilizing a more robust analysis strategy, we were able to identify a total of 15,564 HS-MEs, at an estimated sensitivity and specificity of 98.6% and 99%, respectively. Among these, more than half (8,519) was reported as HS-MEs for the first time, thus achieving a significant improvement in compiling HS-MEs. Such a comprehensive list of HS-MEs provides unique opportunities in examining the patterns of retrotransposition during human genome evolution since the divergence from chimpanzee and gaining new insights regarding the roles of mobile elements in human evolution.

### Challenges in identifying HS-specific MEs

Despite constant improvement in the quality of the reference genome sequences for human and other primates and of the related bioinformatics tools, obtaining a precise list of mobile elements uniquely present in the human genomes faces with many challenges. Several factors contribute to the complications in this task, and these include but are not limited to: 1) incomplete coverage of the reference genome sequences for human and more so for other primates as exemplified in a few recent publications related to human and other primates[39, 58]; 2) assembly errors, particularly in regions rich of MEs; 3) genome rearrangements occurring in a lineage -and species-specific fashion, often involving or mediated by MEs [19–22]; 4) mis- annotation of MEs.

As shown in Table S1, the number of MEs annotated in the human reference genome increased significantly in the most recent version (GRC38, December 2013) compared to an earlier version released in 2004 (GRC35, UCSC hg17)[59], which covered the first major updates since the initial publication of the human draft genome in 2001[1]. The total number of MEs increased ~300,000 (for a 86Mbp increase in non-gap sequences), leading to the increase of ME percentage in the genome from 48.8% in GRC35 to 52.1% in GRC38. The largest increase by ME type is for ERVs, both in terms of the number (366,603) and total sequence size (120.6Mbp), followed by L1, which had an increase of 49,000 in number and 21.8Mbp in size. Interestingly, the change for *Alu* is minimal with the increase in number being ~8,000 and in size being 675Kb. This bias of the degree of improvement, which seems to be correlated with the ME size, is probably due to the fact that the larger the MEs, the more susceptible they are for sequences being incomplete and annotation being incorrect. Apparently, a lot of ERVs were mis-annotated in the earlier versions, and this contributed partly to largest increase in ERVs, and a reduction of “Others” (ME types not specified in Table 1S).

Prior similar studies, best represented by Mills et al[31], were limited by all these factors, as well as by the limitation of methodologies and study scope that either focused on a specific type of MEs or part of the genome[10, 35, 60, 61]. For example, most of the MEs in the repetitive regions, including these in the ME-rich regions, were excluded from all prior studies.

In this study, we tried to address all of these issues. In addition to the use of the best available reference genome sequences for human and other primates, we took an unbiased approach to cover all annotated MEs in the human reference genome, thus representing the first study to include MEs in the repetitive regions, particularly those within other ME rich regions. This has contributed to 3,168 HS-MEs or 37% of the 8,586 novel HS-MEs. Furthermore, we pre-processed the mobile element entries annotated by RepeatMasker to integrate the fragments belong to one original ME but later being interrupted by insertions of other sequences, often by newer MEs or internal rearrangements, such as inversion and deletions. In the case of a full length LTR, RepeatMasker reports three separate entries, representing the two LTR segments and the internal viral sequences. This integration step is important as it restores data to reflect the original number of DNA transposition events and identifies the correct flanking sequences characterizing unique genomic locations of individual MEs. The former corrects the calculation of DNA transposition rate, while the latter is critical for identifying HS-MEs and characterizing their sequences including the TSDs, which is the hallmark of DNA transposition. As shown in Table S1, the fragmentation affects a significant proportion of MEs (as high as 20% in L1s) with the rates seemly to be positively correlated with the element length and the relative ages (data not shown). While impacting mostly older MEs, a total of 845 HS-MEs are also affected, similarly mostly being HS-L1s, likely due to their large insertion size and number of HS-L1 (Table 2.2). The use of multiple non-human primate genomes not only helps reduce the false positive by providing orthologous sequences for gaps or regions of rearrangements in the chimpanzee genome, but also helps increase sensitivity by contributing to 926 novel HS-MEs (Table 2.2).

The use of two sequence alignment tools, blat and liftover, in identifying the orthologous sequences helps reduce the false positives that can be caused by many of the aforementioned complicating factors. Liftover uses alignment data linking closely related genomes through blastz [62], which focuses on large-scale synteny conservation, and is basically the strategy used in the previous analysis of human- and chimp-specific MEs [31]. The method works well for conserved regions, but usually performs poorly for regions involving local rearrangements or with high density of repetitive sequences or misplacement of contigs during assembly. Therefore, it may miss to detect the presence of orthologous MEs and generate high false positives, most like as category II HS-MEs (missing orthologous flanking regions). In contrast, the blat method focuses on identifying local alignments and performs better in handling regions with species-specific sequence changes near MEs, but it may miss some true HS-MEs by generating some false positive detection of orthologous MEs from non-orthologous sequence similarity. Therefore, by requiring the support of both methods for the HS-ME status, we are able to achieve a minimal level of false positive rate in HS-ME detection. We believe that our current list of HS-MEs may represent an underestimate of all HS- MEs that may exist in the human genomes due to the many complications associated with ME analysis and our focus on reducing the false positives.

### The retrotransposition activity level during early human evolution

In the last decade, we and others have extensively studied the profiles of active mobile elements and their relative retrotransposition levels in the human genome mostly based on analysis of polymorphic MEs from personal genomes [10, 32, 33, 36, 46, 47, 52, 61, 63–67]. These studies show that L1s, *Alu*s, SVAs, and HERV-K are the only types of MEs remaining ongoing retrotransposition activity. The most active subfamilies for each ME type have also been established. It is worth to point out that the data reflect the retrotransposition profiles in the current human genomes and during the most recent phase of modern human evolution, and do not reflect the situations during the earlier part of human evolution. The availability of a comprehensive list of ME that uniquely occurred in human in relation to chimpanzee can be used to fill this gap. Based on the assumption that the period for evolution of the *Homo* species is much longer than that for the evolution of modern human populations, we can expect that majority of these HS-MEs occurred during the early phase of human evolution since the separation from chimpanzee. These early HS-MEs would be shared by other archaic human species/subspecies, notably the Neanderthals and Denisovans. Since such data had not been made available before, we can expect to gain new insights about the DNA transposition profiles during the earlier phase of human evolution.

As expected, our data confirmed the previously established most active retrotransposon subfamilies, i.e., Ya5 and Yb8/9 for *Alu*s, L1HS for L1s, the F- subfamilies for SVAs, and HERV-K for LTR retrotransposons (Table 4). In the meantime, notable differences were also observed: the much higher ratio of HS-MEs in relation to all MEs in the same subfamilies and the presence of many extra active subfamilies, including the *Alu*Yf and *Alu*Yk subfamilies from the *Alu* family, the L1P4 and L1M subfamilies from the L1 family, the SVA_A subfamily, and the ERV1 subfamilies from LTR retrotransposons (Table S4). While the high ratios of HS-MEs than that of polymorphic MEs in each active subfamilies may be explained merely by the different time spans, the presence of the extra active subfamilies not seen based on polymorphic MEs may be much better explained by the dropping retrotransposition activity in the modern human genomes.

### Y chromosome as a hot target for HS-LTRs

As shown in Table S6 and Fig. 2, the HS-ME density on the Y chromosome is significantly higher than all other chromosomes. By ME type, *Alu*, L1, and LTR MEs all show higher density for Y chromosome. For HS-*Alu*s and HS-L1s, the density in Y chromosome is at least 2 times higher than the genome average, while for HS-LTRs, the density in Y chromosome is close to 40 times higher than the genome average. This is really highly unusual and unexpected to us. This extreme bias for Y chromosome cannot seem to be explained by any factors that we are aware for impacting the ME distribution in the genome, such as gene density, GC content, poorer sequence quality. For gene density, chromosome 13 is similarly low in gene density as Y chromosome, but its HS-LTR density is in fact higher than many other autosomes that have higher gene density (Table S6). Contrary to HS-LTRs, HS-SVAs showed a strong bias against Y chromosome with the density being ~10% of the genome average, which can be partially explained by HS-SVAs’ strong preference for gene rich regions (Pearson coefficience=0.9 with gene density). However, the density of HS-SVAs in chromosome 13, which has a similar gene density as Y chromosome, is close to the genome average and is more than 6 times higher than of Y chromosome. If such bias is caused by lower sequence quality (more sequence gaps) in both human and chimpanzee for Y chromosome and less sequence available for non-primates, we would expect to see a similar trend for all types of MEs, and not see an opposite pattern between SVAs and LTRs.

**Figure 3.**
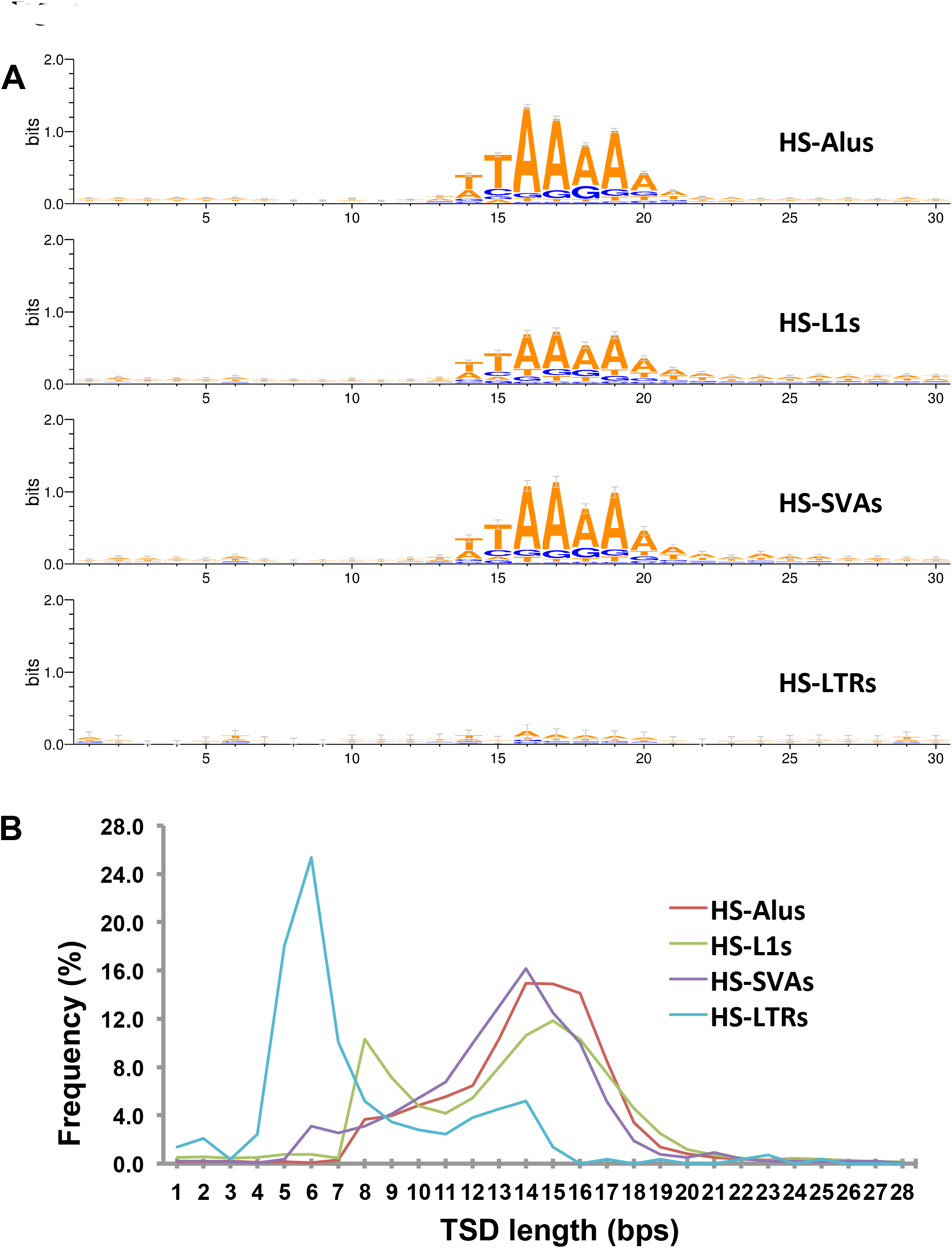
Characteristics of pre-integration sites and TSD lengths: A. Sequence logo for the pre-integration sites; B. TSD length distributions

According to the male germline hypothesis, the densities of MEs on each chromosome should positively correlate with the amount of time spent in the male germline. Assuming cell division is mutagenic and the number of germ cell divisions is larger in males than in females (as is likely in most mammals), male is expected to be main source of mutations. The autosomes have a ½ chance carried by the male while chromosome Y is always carried by the male. Chromosome X, however, has a 1/3 chance carried by the male. In theory, as per generation, the maximum mutation frequency between autosomes and sex chromosomes would be Y:A:X = 6:3:2 [57]. This hypothesis is supported by the observed Y>A>X by density of young *Alu*s [68].

Another hypothesis is that the ME density on Y chromosome is facilitated by the low recombination-driven deletion in chromosome Y. As Y chromosome does not recombine with chromosome X outside of the pseudo-autosomal regions, it would have higher ME density than the autosomes. However, this hypothesis is not supported by our results that the HS-RE density and shared RE density are higher on the pseudo- autosomal regions than the male specific regions on chromosome Y (data not shown).

Natural selection might also be contributing to the Y chromosome bias: Y chromosome has a lower frequency of DNA recombination due to lack of homologous regions [69]. DNA recombination-mediated deletion can serve as a mechanism for removing ME insertions in a way biased for those with negative functional impact via selection. Because Y chromosome has much fewer genes than any other chromosomes, it is facing a lower level of selection pressure and lower rate of insertion removal.

However, none of these mentioned hypotheses could explain why this bias is observed for HS-MEs but not for older ME that are shared with other primates, nor can they explain why HS-SVAs is showing the opposite. Although genetic decay may be the principal dynamic in the evolution of chromosome Y, recent studies suggested that remodeling and regeneration have dominated chimpanzee and human male specific Y chromosome evolution during the past 6 million years [48]. This could potentially explain the success of HS-MEs on chromosome Y.

Assuming this bias is valid, how would it affect chromosome Y? Recently, there has been a heated debate among the science community about whether chromosome Y is disappearing or not [70]. Our result could serve as evidence that “chromosome Y has not disappeared yet” and HS-MEs may have contributed to the observed fast evolving pattern on chromosome Y after the human and chimpanzee divergence [48]. However, the potential functional impact about the preferential insertion of HS-MEs, particularly the LTR retrotransposons on Y chromosome is still unknown.

### The impact of MEs on human genome evolution and gene function

The 15,586 HS-MEs collectively contribute a net increase of the genome size by 14.5 million base pairs. This may be the only defined significant change in the human genome during human evolution. This size change is significant in a relatively short period of time by a single molecular mechanism by way of retrotransposition. This size of genome increase is close to one quarter of the Y chromosome and is larger than the genomes of free-living eukaryotic organisms, such as yeast [71]. The only other molecular mechanism that could contribute a significant size change to the human genome would be genomic segmental duplication [72]. However, no data is available about the exact amount of human-specific segmental duplication occurred for the same period of human evolution.

Among the HS-MEs, a total of 8,001 or 52% of HS-MEs are located inside or in the 1kb promoter regions of genes for protein coding, non-coding RNAs, as well as transcribed pseudogenes, representing 4,713 unique genes/transcripts (Table 4). In 284 cases, these HS-MEs are part of transcripts, representing mostly alternative splice forms. Interestingly, in 7 of these cases a HS-ME contribute to part of the coding region in the transcripts, albeit all being rare splice form documented the GENCODE transcripts, not in NCBI RefGene list. Furthermore, 1,213 of the HS-MEs contribute to 3,124 binding sites for 146 of the 161 examined transcriptional factors in association with 622 genes (data not shown). For those in the intron regions that represent most of the 8,001 HS-MEs in genic regions, while a direct functional impact would be hard to predict, they may have potential in participating in regulation of transcription and alternative splicing as demonstrated by documented examples[16, 23].

In summary, we data suggest that, despite being very young in the genome, many of these HS-MEs already have already participated in gene function via regulation of transcription, splicing, and protein coding, and there may be more potential for their future participation as demonstrated by Ward et al [18, 28].

### Future directions

Due to the technical challenges associated with the analysis of mobile elements and deficiencies of reference genome sequences for human and other primates, our list of HS-MEs still suffers a certain level of false negatives and false positives. We can expect that the number of HS-MEs continue to increase from regions with sequencing gaps, especially regions in highly rich of repetitive sequences, such as the centromere and telomere regions, which may be hot spots for certain types of ME insertion, such as LTRs [73]. While the current list of HS-MEs covers those appeared in the human genome after human and chimpanzee separation, we can further divide these HS-MEs to those unique to *Homo sapiens* vs. those in all *Homo* species by comparative analysis of such genome sequences (e.g. the neanderthal and Denisovan genomes) [74]. An additional note is that since the identification of HS-MEs is based on a consensus genome sequence, which is mostly of a Caucasian origin, some of these MEs are polymorphic being present in some but not all humans. In the meantime, we are missing the non-reference MEs. In the future, it should be useful to generate a list of HS-MEs common to all humans (minimal set of MEs) and another list of HS-MEs that are polymorphic. The former would be useful for analyzing MEs’ impact on evolution of all modern humans, while the latter would be useful for studying MEs’ contribution to genetic and phenotypic diversity among human populations and individuals.

To better understand the potential functional impact of these HS-MEs, we may be able to use transcriptome data, as well as epigenetic and other functional genomic data, such as genome-wide DNA methylation, histone modification profiling data, and transcroptome data for transcriptional factors, with representation of individuals from diverse population backgrounds and for all human tissues. These data that are being accumulated rapidly will help us better assess the role of these HS-MEs in generating new regulatory elements, alternative splicing, novel transcripts, and epigenetic regulation.

## Acknowledgements

This work is in part supported by grants from the Canadian Research Chair program, Canadian Foundation of Innovation, Ontario Ministry of Research and Innovation, Canadian Natural Science and Engineering Research Council, and Brock University to PL, and was made possible by Compute Canada high performance computing facilities.

## List of tables

**Table 1: A summary of MEI counts and HS-MEIs.**

**Table 2: Sources of novel HS-MEIs.**

**Table 3: Impact of HS-MEIs to genome size**

**Table 4: Distribution of HS-MEIs by gene context**

## List of supplementary Figures

**Figure S1. Selective representative of PCR verification**

PCR result for two cases of HS-MEIs in MEIs and HS-MEIs in HS-MEIs.

**Figure S2. Examples of HS-MEIs from different sources.**

A: false negative using chimp sequence only
B: false negative due to MEI fragmentation
c: false negative in Mills et al, 2006
D: HS-MEI in HS-MEI.

**Figure S3. GC region of HS-MEI and shared-MEI**

## List of supplementary tables

**Table S1: Mobile element compositions in different versions of the human reference genome**

[Based on Table S1, provide a revised counting for the number of MEIs and their exact percentage of size contribution to the genome. Breaking down to main types, SINE, LINE, LTR, Transposons, Others; under each main type, list the top 1 or two main subtypes, e.g.: *Alu*, L1, SVA, and HERV; with their original counts based on RepeatMaskers and adjusted counts, size contribution: insertion only and total (including TSD, transductions)]

**Table S2. Comparison of HS-MEIs with results from prior studies.**

Columns: MEI type, Mills et al 2009, Tang et al 2016, shared, uniqToMills2009, uniq2Tang2016”;

Rows: *Alu*, L1, SVA, LTR, and Total.

**Table S3. Result of PCR validation**

Columns: HS-MEI_ID, location, MEI-in-MEI type (e.g. *Alu* in SVA), primers (5’, 3’), HS- MEI status

**Table S4. Retrotransposition activities of MEIs from different subfamilies. Table S5. Distribution of MEIs-in-MEIs**

**Table S6. Distributions of HS-MEIs by chromosome**

## List of supplementary files

**SupplementaryFile1: Supplementary information on materials and methods Supplementary**

**File2: A complete HS-MEI list**

[In tab delimited text file, provide the following items of information for all HS-MEIs: HS-MEI_ID (use dbRIP IDs); the genome coordinates (hg19), family and subfamily (three levels); length (excluding MEI-in-MEIs), TSD: length, sequence or NA), Category (I or II).

## References

1. Lander ES, Linton LM, Birren B, Nusbaum C, Zody MC, Baldwin J, Devon K, Dewar K, Doyle M, FitzHugh W et al: Initial sequencing and analysis of the human genome. Nature 2001, 409(6822):860–921.

2. Deininger PL, Moran JV, Batzer MA, Kazazian HH Jr.: Mobile elements and mammalian genome evolution. Curr Opin Genet Dev 2003, 13(6):651–658.

3. Cordaux R, Batzer MA: The impact of retrotransposons on human genome evolution. Nat Rev Genet 2009, 10(10):691–703.

4. Kazazian HH Jr.: Mobile elements: drivers of genome evolution. Science 2004, 303(5664):1626–1632.

5. Ostertag EM, Kazazian HH Jr.: Biology of mammalian L1 retrotransposons. Annu Rev Genet 2001, 35:501–538.

6. Batzer MA, Deininger PL: Alu repeats and human genomic diversity. Nat Rev Genet 2002, 3(5):370–379.

7. Kazazian HH Jr., Moran JV: The impact of L1 retrotransposons on the human genome. Nat Genet 1998, 19(1):19–24.

8. Kimberland ML, Divoky V, Prchal J, Schwahn U, Berger W, Kazazian HH Jr.: Full-length human L1 insertions retain the capacity for high frequency retrotransposition in cultured cells. Hum Mol Genet 1999, 8(8):1557–1560.

9. Kazazian HH Jr., Goodier JL: LINE drive. retrotransposition and genome instability. Cell 2002, 110(3):277–280.

10. Mills RE, Bennett EA, Iskow RC, Devine SE: Which transposable elements are active in the human genome? Trends Genet 2007, 23(4):183–191.

11. Ostertag EM, Goodier JL, Zhang Y, Kazazian HH Jr.: SVA elements are nonautonomous retrotransposons that cause disease in humans. Am J Hum Genet 2003, 73(6):1444–1451.

12. Wang H, Xing J, Grover D, Hedges DJ, Han K, Walker JA, Batzer MA: SVA elements: a hominid-specific retroposon family. J Mol Biol 2005, 354(4):994–1007.

13. Doolittle WF, Sapienza C: Selfish genes, the phenotype paradigm and genome evolution. Nature 1980, 284(5757):601–603.

14. Symer DE, Connelly C, Szak ST, Caputo EM, Cost GJ, Parmigiani G, Boeke JD: Human l1 retrotransposition is associated with genetic instability in vivo. Cell 2002, 110(3):327–338.

15. Szak ST, Pickeral OK, Landsman D, Boeke JD: Identifying related L1 retrotransposons by analyzing 3' transduced sequences. Genome Biol 2003, 4(5):R30.

16. Han JS, Szak ST, Boeke JD: Transcriptional disruption by the L1 retrotransposon and implications for mammalian transcriptomes. Nature 2004, 429(6989):268- 274.

17. Wheelan SJ, Aizawa Y, Han JS, Boeke JD: Gene-breaking: a new paradigm for human retrotransposon-mediated gene evolution. Genome Res 2005, 15(8):1073–1078.

18. Mita P, Boeke JD: How retrotransposons shape genome regulation. Curr Opin Genet Dev 2016, 37:90–100.

19. Callinan PA, Wang J, Herke SW, Garber RK, Liang P, Batzer MA: Alu retrotransposition-mediated deletion. J Mol Biol 2005, 348(4):791–800.

20. Han K, Sen SK, Wang J, Callinan PA, Lee J, Cordaux R, Liang P, Batzer MA: Genomic rearrangements by LINE-1 insertion-mediated deletion in the human and chimpanzee lineages. Nucleic Acids Res 2005, 33(13):4040–4052.

21. Sen SK, Han K, Wang J, Lee J, Wang H, Callinan PA, Dyer M, Cordaux R, Liang P, Batzer MA: Human genomic deletions mediated by recombination between Alu elements. Am J Hum Genet 2006, 79(1):41–53.

22. Han K, Lee J, Meyer TJ, Wang J, Sen SK, Srikanta D, Liang P, Batzer MA: Alu recombination-mediated structural deletions in the chimpanzee genome. PLoS Genet 2007, 3(10):1939–1949.

23. Quinn JP, Bubb VJ: SVA retrotransposons as modulators of gene expression. Mob Genet Elements 2014, 4:e32102.

24. Konkel MK, Batzer MA: A mobile threat to genome stability: The impact of non-LTR retrotransposons upon the human genome. Semin Cancer Biol 2010, 20(4):211–221.

25. Hancks DC, Kazazian HH Jr.: Active human retrotransposons: variation and disease. Curr Opin Genet Dev 2012, 22(3):191–203.

26. Callinan PA, Batzer MA: Retrotransposable elements and human disease. Genome Dyn 2006, 1:104–115.

27. Ahmed M, Liang P: Transposable elements are a significant contributor to tandem repeats in the human genome. Comp Funct Genomics 2012, 2012:947089.

28. Ward MC, Wilson MD, Barbosa-Morais NL, Schmidt D, Stark R, Pan Q, Schwalie PC, Menon S, Lukk M, Watt S et al: Latent regulatory potential of human-specific repetitive elements. Mol Cell 2013, 49(2):262–272.

29. Chuong EB, Elde NC, Feschotte C: Regulatory evolution of innate immunity through co-option of endogenous retroviruses. Science 2016, 351(6277):1083- 1087.

30. Huang CR, Schneider AM, Lu Y, Niranjan T, Shen P, Robinson MA, Steranka JP, Valle D, Civin CI, Wang T et al: Mobile interspersed repeats are major structural variants in the human genome. Cell 2010, 141(7):1171–1182.

31. Mills RE, Bennett EA, Iskow RC, Luttig CT, Tsui C, Pittard WS, Devine SE: Recently mobilized transposons in the human and chimpanzee genomes. Am J Hum Genet 2006, 78(4):671–679.

32. Wang J, Song L, Gonder MK, Azrak S, Ray DA, Batzer MA, Tishkoff SA, Liang P: Whole genome computational comparative genomics: A fruitful approach for ascertaining Alu insertion polymorphisms. Gene 2006, 365:11–20.

33. Ahmed M, Li W, Liang P: Identification of three new Alu Yb subfamilies by source tracking of recently integrated Alu Yb elements. Mob DNA 2013, 4(1):25.

34. Hughes JF, Coffin JM: Human endogenous retrovirus K solo-LTR formation and insertional polymorphisms: implications for human and viral evolution. Proc Natl Acad Sci U S A 2004, 101(6):1668–1672.

35. Buzdin A, Ustyugova S, Khodosevich K, Mamedov I, Lebedev Y, Hunsmann G, Sverdlov E: Human-specific subfamilies of HERV-K (HML-2) long terminal repeats: three master genes were active simultaneously during branching of hominoid lineages. Genomics 2003, 81(2):149–156.

36. Shin W, Lee J, Son SY, Ahn K, Kim HS, Han K: Human-specific HERV-K insertion causes genomic variations in the human genome. PLoS One 2013, 8(4):e60605.

37. Chimpanzee S, Analysis C: Initial sequence of the chimpanzee genome and comparison with the human genome. Nature 2005, 437(7055):69–87.

38. Scally A, Dutheil JY, Hillier LW, Jordan GE, Goodhead I, Herrero J, Hobolth A, Lappalainen T, Mailund T, Marques-Bonet T et al: Insights into hominid evolution from the gorilla genome sequence. Nature 2012, 483(7388):169–175.

39. Gordon D, Huddleston J, Chaisson MJ, Hill CM, Kronenberg ZN, Munson KM, Malig M, Raja A, Fiddes I, Hillier LW et al: Long-read sequence assembly of the gorilla genome. Science 2016, 352(6281):aae0344.

40. Locke DP, Hillier LW, Warren WC, Worley KC, Nazareth LV, Muzny DM, Yang SP, Wang Z, Chinwalla AT, Minx P et al: Comparative and demographic analysis of orang-utan genomes. Nature 2011, 469(7331):529–533.

41. Carbone L, Harris RA, Gnerre S, Veeramah KR, Lorente-Galdos B, Huddleston J, Meyer TJ, Herrero J, Roos C, Aken B et al: Gibbon genome and the fast karyotype evolution of small apes. Nature 2014, 513(7517):195–201.

42. Rhesus Macaque Genome S, Analysis C, Gibbs RA, Rogers J, Katze MG, Bumgarner R, Weinstock GM, Mardis ER, Remington KA, Strausberg RL et al: Evolutionary and biomedical insights from the rhesus macaque genome. Science 2007, 316(5822):222–234.

43. Zimin AV, Cornish AS, Maudhoo MD, Gibbs RM, Zhang X, Pandey S, Meehan DT, Wipfler K, Bosinger SE, Johnson ZP et al: A new rhesus macaque assembly and annotation for next-generation sequencing analyses. Biol Direct 2014, 9(1):20.

44. Yan G, Zhang G, Fang X, Zhang Y, Li C, Ling F, Cooper DN, Li Q, Li Y, van Gool AJ et al: Genome sequencing and comparison of two nonhuman primate animal models, the cynomolgus and Chinese rhesus macaques. Nat Biotechnol 2011, 29(11):1019–1023.

45. Kent WJ: BLAT–the BLAST-like alignment tool. Genome Res 2002, 12(4):656–664.

46. Stewart C, Kural D, Stromberg MP, Walker JA, Konkel MK, Stutz AM, Urban AE, Grubert F, Lam HY, Lee WP et al: A comprehensive map of mobile element insertion polymorphisms in humans. PLoS Genet 2011, 7(8):e1002236.

47. Sudmant PH, Rausch T, Gardner EJ, Handsaker RE, Abyzov A, Huddleston J, Zhang Y, Ye K, Jun G, Hsi-Yang Fritz M et al: An integrated map of structural variation in 2,504 human genomes. Nature 2015, 526(7571):75–81.

48. Hughes JF, Skaletsky H, Pyntikova T, Graves TA, van Daalen SK, Minx PJ, Fulton RS, McGrath SD, Locke DP, Friedman C et al: Chimpanzee and human Y chromosomes are remarkably divergent in structure and gene content. Nature 2010, 463(7280):536–539.

49. Harrow J, Frankish A, Gonzalez JM, Tapanari E, Diekhans M, Kokocinski F, Aken BL, Barrell D, Zadissa A, Searle S et al: GENCODE: the reference human genome annotation for The ENCODE Project. Genome Res 2012, 22(9):1760–1774.

50. Pruitt KD, Tatusova T, Maglott DR: NCBI reference sequences (RefSeq): a curated non-redundant sequence database of genomes, transcripts and proteins. Nucleic Acids Res 2007, 35(Database issue):D61–65.

51. Mills RE, Luttig CT, Larkins CE, Beauchamp A, Tsui C, Pittard WS, Devine SE: An initial map of insertion and deletion (INDEL) variation in the human genome. Genome Res 2006, 16(9):1182–1190.

52. Wang J, Song L, Grover D, Azrak S, Batzer MA, Liang P: dbRIP: a highly integrated database of retrotransposon insertion polymorphisms in humans. Hum Mutat 2006, 27(4):323–329.

53. Abrusan G, Krambeck HJ: The distribution of L1 and Alu retroelements in relation to GC content on human sex chromosomes is consistent with the ectopic recombination model. J Mol Evol 2006, 63(4):484–492.

54. Jurka J: Sequence patterns indicate an enzymatic involvement in integration of mammalian retroposons. Proc Natl Acad Sci U S A 1997, 94(5):1872–1877.

55. Cost GJ, Boeke JD: Targeting of human retrotransposon integration is directed by the specificity of the L1 endonuclease for regions of unusual DNA structure. Biochemistry 1998, 37(51):18081–18093.

56. Bushman FD: Targeting survival: integration site selection by retroviruses and LTR-retrotransposons. Cell 2003, 115(2):135–138.

57. Genomes Project C, Abecasis GR, Altshuler D, Auton A, Brooks LD, Durbin RM, Gibbs RA, Hurles ME, McVean GA: A map of human genome variation from population-scale sequencing. Nature 2010, 467(7319):1061–1073.

58. Kim JI, Ju YS, Park H, Kim S, Lee S, Yi JH, Mudge J, Miller NA, Hong D, Bell CJ et al: A highly annotated whole-genome sequence of a Korean individual. Nature 2009, 460(7258):1011–1015.

59. International Human Genome Sequencing C: Finishing the euchromatic sequence of the human genome. Nature 2004, 431(7011):931–945.

60. Barbulescu M, Turner G, Seaman MI, Deinard AS, Kidd KK, Lenz J: Many human endogenous retrovirus K (HERV-K) proviruses are unique to humans. Curr Biol 1999, 9(16):861–868.

61. Bennett EA, Keller H, Mills RE, Schmidt S, Moran JV, Weichenrieder O, Devine SE: Active Alu retrotransposons in the human genome. Genome Res 2008, 18(12):1875–1883.

62. Schwartz S, Kent WJ, Smit A, Zhang Z, Baertsch R, Hardison RC, Haussler D, Miller W: Human-mouse alignments with BLASTZ. Genome Res 2003, 13(1):103–107.

63. Konkel MK, Wang J, Liang P, Batzer MA: Identification and characterization of novel polymorphic LINE-1 insertions through comparison of two human genome sequence assemblies. Gene 2007, 390(1-2):28–38.

64. Lee J, Cordaux R, Han K, Wang J, Hedges DJ, Liang P, Batzer MA: Different evolutionary fates of recently integrated human and chimpanzee LINE-1 retrotransposons. Gene 2007, 390(1-2):18–27.

65. Wildschutte JH, Williams ZH, Montesion M, Subramanian RP, Kidd JM, Coffin JM: Discovery of unfixed endogenous retrovirus insertions in diverse human populations. Proc Natl Acad Sci U S A 2016, 113(16):E2326–2334.

66. Iskow RC, McCabe MT, Mills RE, Torene S, Pittard WS, Neuwald AF, Van Meir EG, Vertino PM, Devine SE: Natural mutagenesis of human genomes by endogenous retrotransposons. Cell 2010, 141(7):1253–1261.

67. Witherspoon DJ, Zhang Y, Xing J, Watkins WS, Ha H, Batzer MA, Jorde LB: Mobile element scanning (ME-Scan) identifies thousands of novel Alu insertions in diverse human populations. Genome Res 2013, 23(7):1170–1181.

68. Jurka J, Krnjajic M, Kapitonov VV, Stenger JE, Kokhanyy O: Active Alu elements are passed primarily through paternal germlines. Theor Popul Biol 2002, 61(4):519- 530.

69. Boissinot S, Entezam A, Furano AV: Selection against deleterious LINE-1- containing loci in the human lineage. Mol Biol Evol 2001, 18(6):926–935.

70. Griffin DK: Is the Y chromosome disappearing?–both sides of the argument. Chromosome Res 2012, 20(1):35–45.

71. Cherry JM, Ball C, Weng S, Juvik G, Schmidt R, Adler C, Dunn B, Dwight S, Riles L, Mortimer RK et al: Genetic and physical maps of Saccharomyces cerevisiae. Nature 1997, 387(6632 Suppl):67–73.

72. Bailey JA, Eichler EE: Primate segmental duplications: crucibles of evolution, diversity and disease. Nat Rev Genet 2006, 7(7):552–564.

73. Zahn J, Kaplan MH, Fischer S, Dai M, Meng F, Saha AK, Cervantes P, Chan SM, Dube D, Omenn GS et al: Expansion of a novel endogenous retrovirus throughout the pericentromeres of modern humans. Genome Biol 2015, 16:74.

74. Ahmed M, Liang P: Study of Modern Human Evolution via Comparative Analysis with the Neanderthal Genome. Genomics Inform 2013, 11(4):230–238.

